# Longer study length, standardized sampling techniques, and broader geographic scope leads to higher likelihood of detecting stable abundance patterns in long term black-legged tick studies

**DOI:** 10.1101/2021.03.06.434217

**Authors:** Rowan Christie, Kaitlin Stack Whitney, Julia Perrone, Christine A. Bahlai

## Abstract

**Background:** Understanding how study design and monitoring strategies shape inference within, and synthesis across, studies is critical across biological disciplines. Many biological and field studies are short term and limited in scope. Monitoring studies are critical for informing public health about potential vectors of concern, such as *Ixodes scapularis* (black-legged ticks). Black-legged ticks are a taxon of ecological and human health concern due to their status as primary vectors of *Borrelia burgdorferi*, which causes Lyme disease. However, variation in black-legged tick monitoring, and gaps in data, are currently considered major barriers to understanding population trends and in turn, predicting Lyme disease risk. To understand how variable methodology in black-legged tick studies may influence which population patterns researchers find, we conducted a data synthesis experiment.

**Materials and Methods:** We searched for publicly available black-legged tick abundance datasets that had at least 9 years of data, using keywords about ticks in internet search engines, literature databases, data repositories and public health websites. Our analysis included 289 datasets from 7 surveys from locations throughout the US, ranging in length from 9 to 24 years. We used a moving window analysis, which is a kind of non-random resampling approach, to investigate the temporal stability of black-legged tick population trajectories across the US. We then used t-tests to assess differences in stability time across different study parameters.

**Results:** All of our sampled datasets required 4 or more years to reach stability. We also found that several study factors can have an impact on the likelihood of a study reaching stability and of data resulting in misleading results if the study does not reach stability. Specifically, datasets collected via dragging reached stability significantly faster than data collected via opportunistic sampling. Datasets that sampled larva reached stability significantly later than those that sampled adults or nymphs. Additionally, datasets collected at the broadest spatial scale (county) reached stability fastest.

**Conclusion:** We used 289 datasets from 7 long term black-legged tick studies to conduct a non-random data resampling experiment, revealing that sampling design does shape inferences in black-legged tick population trajectories and how many years it takes to find stable patterns. Specifically, our results show the importance of study length, sampling technique, life stage, and geographic scope in understanding black-legged tick populations, in the absence of standardized surveillance methods. Current public health efforts based on existing black-legged tick datasets must take monitoring study parameters into account, to better understand if and how to use monitoring data to inform decision making. We also recommend that potential forecasting initiatives consider these parameters when projecting black-legged tick population trends.

## Introduction

Understanding how study design and monitoring strategies shape inference within, and synthesis across, studies is critical across biological sciences. This is especially important as climatic conditions and ecosystems change in stark and unpredictable ways, requiring scientists to tease apart drivers of population and community trends from the past to understand possible futures (Bahlai et al., 2021). Many biological and field studies are short term and limited in scope, leading to concerns about how to interpret the many short-term studies in comparison with the relatively fewer long term ecological studies (Wauchope et al., 2019; Bahlai et al., 2021). Shorter term studies increase the likelihood of finding misleading trends, potentially leading to misinformed management approaches (White and Bahlai, 2021). Natural variability in a stable population might lead a 2-3 year study to appear to capture a strong upward trend in abundance for a given taxon, simply as a result of natural population cycling (Fournier et al., 2019).

Monitoring studies are critical for informing public health about potential vectors of concern. Surveys indicate that the control and management of vector borne diseases is of immediate concern for public health in the United States (Hill et al., 2009). Public health officials must know the geographical range of infectious vectors and their level of abundance to understand potential disease risk (Townson et al., 2005). However, standardizing results from multiple studies on vectors can be problematic. Novel sampling approaches and methodologies are required to effectively monitor variable vector populations (Sougoufara et al., 2020). Reliable inference about population trends from single studies, or synthesizing across them, must account for variation in monitoring methods.

A critical vector of concern for public health in the US is ticks. They are common vectors that can transmit multiple diseases to humans and are responsible for vectoring the greatest number of human diseases in the US (Eisen et al., 2017). Increased outdoor activity, especially during the ongoing COVID-19 pandemic, may be exposing more people to potential tick-borne illnesses (HHS 2020). This pattern matches broader tick trends. Globally, both the known range of tick species and the reported cases of tick-vectored illnesses are increasing (Paddock et al., 2016).

Ticks and tick-vectored diseases are commonly monitored using dragging, flagging, and carbon dioxide baited traps (Falco and Fish, 1992). Dragging is a method where a person pulls a horizontal pole behind them across the ground and ticks latch onto the cloth attached to the pole (Salomon et al., 2020). Another common tick survey method is passive surveillance, where individuals send in ticks they have found on themselves to public health departments, but do not deliberately search for ticks (Ripoche et al., 2018). While these are common, there is a wide variety of methods reported across studies. For example, one method of surveying ticks has been collecting them from hunter-killed deer in the midwestern US (Egizi et al., 2018; Lee et al., 2013). Additionally, some studies measure density instead of abundance (Kahl et al., 2002), use distance-based sampling (Gray et al., 1992), or are modeled projections of where ticks are likely to be (Baldwin et al., 2022), based on varying assumptions about climate and landscape factors (Estrada-Peña et al., 2013). The lack of a shared and standardized sampling method may be preventing effective understanding of changes in abundance and distribution of ticks synthesized across surveys (Dennis et al., 1998; Eisen et al., 2016).

The lack of a standardized sampling method may also impact the ability to accurately monitor tick-vectored disease. Recent research has identified the lack of standardized data collection, and subsequent data sharing, as impeding researchers’ understanding of tick-transmitted pathogens (Estrada-Peña et al., 2021).

This is critical, as Lyme disease is a common and increasing illness impacting US residents. Recent research estimates that more than 400,000 people annually have been diagnosed in the US between 2010 and 2018 (Kugeler et al., 2021). The primary vectors of *Borrelia burgdorferi*, bacteria that cause Lyme disease, are Black-legged ticks (Des Vignes and Fish, 1997). Black-legged ticks are common throughout the US - and both widespread and abundant. Established black-legged tick populations are most concentrated in upper north-central, northeastern, and west-coast states (Eisen et al. 2016). Black-legged ticks transmit seven human pathogens of concern, including *B. burgdorferi*, and both the tick and pathogen transmission have increased over the past two decades in the US (Eisen and Eisen 2018). In addition, Black-legged ticks are spreading rapidly, found in more than twice the number of counties in the United States compared to twenty years ago (Eisen et al. 2016). Systematic surveillance of ticks and tick-borne pathogens is important for understanding Lyme disease risk (Eisen and Paddock 2021). Variation in black-legged tick monitoring, and gaps in data, are currently considered major barriers to predicting Lyme disease risk (Kugeler and Eisen 2020). Understanding tick prevalence and potential Lyme disease risk can help shift the burden of tick vectored disease management from individual prevention to states or other public health agencies (Eisen 2020).

To understand how variable methodology in black-legged tick studies may influence which population patterns researchers find, we conducted a data synthesis experiment. To do this, we used a novel nonrandom sampling approach (Bahlai et al., 2021) to analyze publicly available black-legged tick datasets from long term monitoring studies to understand the impacts of study parameters on inferences about black-legged tick abundance and trajectory. Specifically, we examined study length, sampling technique, life stage, geographic scope, and sampling metrics. We anticipated that studies using dragging would have less variable findings than other sampling techniques, due to existing research showing that dragging is more consistent for some life stages of black-legged tick collection (Falco and Fish, 1992). We also anticipated that the tick life stage monitored could influence expected trends. Nymphs are most likely to spread *Borrelia burgdorferi* (Rulison et al., 2013), and may have different abundance patterns than adults or larvae. Results from this analysis can help researchers contextualize existing monitoring data and the potential opportunities and challenges when extrapolating from the past to the future. This data experiment may also help researchers better understand how to design black-legged tick monitoring studies to avoid misleading inferences, potentially leading to improved surveillance of a public health risk.

## Materials & Methods

### Data searching

We searched for publicly available black-legged tick abundance dataset that had at least 9 years of data, using keywords about black-legged ticks in internet search engines, literature databases, data repositories and public health websites. Specifically, we searched combinations of *tick* and *scapularis* in Data Dryad, Data One, Google Datasets, the Long Term Ecological Research data portal, and the National Ecological Observation Network data portal. We then used Google search portals to look up individual state department of health websites and state datasets where black-legged ticks are common for additional sets of survey data. Our complete search was done using the PRISMA workflow for systematic literature screening, review and reporting (Moher et al., 2009; Appendix 1).

### Black-legged tick datasets

Our search yielded 289 datasets from 7 surveys from locations in New Jersey, New York, Iowa, Connecticut, Massachusetts, ranging in length from 9 years to 24 years. We extracted and scored information about each dataset in terms of the life stage sampled, sampling techniques used, geographic scale of the study, and location (Table 1). The surveys varied in the life stages included (Table 1). Black-legged ticks go through four life stages: egg, larva, nymph, and adult (CDC 2011). Our search yielded surveys that collected data on adults only, nymphs only, adults and nymphs, or all life stages other than egg. These included datasets that studied adults (oldest life stage, n=63 datasets), nymphs (middle life stage, n=68 datasets), and larvae (youngest life stage, n=8 datasets). Additional datasets (n=150) did not specify the life stage collected and were not included in subsequent analyses about life stage. The search results also included datasets that either only included abundance - or also included analysis of the percentage of ticks infected with *B. burgdorferi*. Across the surveys, we found most measured the abundance of black-legged ticks (n=175 datasets) and some tested which percentage had *B. burgdorferi* (n=114 datasets). Surveys also varied in their geographic scope (Table 1). Three surveys were collected and reported at the county scale, while two were reported by town, and two at smaller spatial scales (grid and state forest). The studies included datasets summarized by grid (n=24 datasets), state forest (n=6 datasets), town (n=186 datasets), and county (n=73 datasets). The surveys found in our search also included two sampling techniques, dragging (n=90 datasets) and opportunistic surveying of ticks found on individuals (n=198 datasets) (Table 1). One additional dataset instead recorded “bites found on a person” (n=1), instead of ticks, and was not included in subsequent analyses.

**Table 1.**
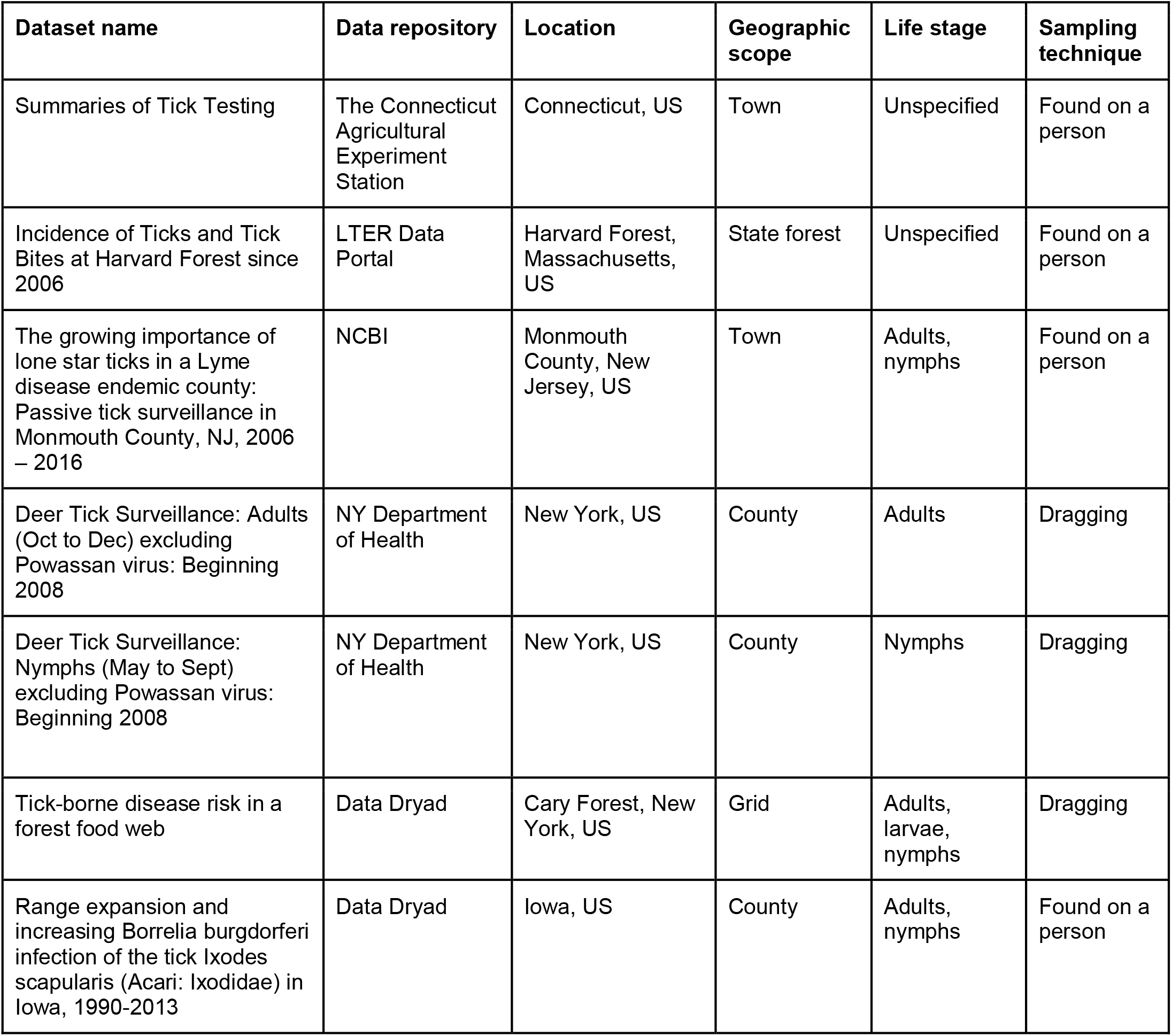
Location, geographic scope, life stage, and sampling technique of black-legged deer tick studies found that met our study criteria (Appendix 1). Datasets from these studies were included in our analyses.

| Dataset name | Data repository | Location | Geographic scope | Life stage | Sampling technique |
| --- | --- | --- | --- | --- | --- |
| Summaries of Tick Testing | The Connecticut Agricultural Experiment Station | Connecticut, US | Town | Unspecified | Found on a person |
| Incidence of Ticks and Tick Bites at Harvard Forest since 2006 | LTER Data Portal | Harvard Forest, Massachusetts, US | State forest | Unspecified | Found on a person |
| The growing importance of lone star ticks in a Lyme disease endemic county: Passive tick surveillance in Monmouth County, NJ, 2006 – 2016 | NCBI | Monmouth County, New Jersey, US | Town | Adults, nymphs | Found on a person |
| Deer Tick Surveillance: Adults (Oct to Dec) excluding Powassan virus: Beginning 2008 | NY Department of Health | New York, US | County | Adults | Dragging |
| Deer Tick Surveillance: Nymphs (May to Sept) excluding Powassan virus: Beginning 2008 | NY Department of Health | New York, US | County | Nymphs | Dragging |
| Tick-borne disease risk in a forest food web | Data Dryad | Cary Forest, New York, US | Grid | Adults, larvae, nymphs | Dragging |
| Range expansion and increasing *Borrelia burgdorferi* infection of the tick *Ixodes scapularis* (Acari: Ixodidae) in Iowa, 1990-2013 | Data Dryad | Iowa, US | County | Adults, nymphs | Found on a person |

### Moving window analysis

To analyze the stability of the abundance trends of these datasets, we used the ‘broken window’ algorithm (Bahlai et al. 2021) using RStudio and R statistical software (RStudio Team, 2020; R Core Team, 2020). This non-random resampling approach uses subsamples (i.e. ‘*windows*’) of the time series data to gain insights into patterns of how data behaves in arbitrarily selected time periods (Bahlai et al. 2021). Non-random resampling of existing monitoring data is a powerful and underused tool to understand trends (White and Bahlai 2021).

This algorithm defines the linear slope associated with the longest time series available as the ‘true’ slope (as it is calculated using the largest possible sample size) and performs calculations relative to this proxy for comparison. The algorithm uses a yearly interval for its calculations, and in the case of this study, the metric used as the dependent variable was some unit of abundance or activity of ticks, as defined by the trapping method. Prior to subsetting within the algorithm, each raw dataset is subjected to normalization by converting it to a unitless Z score, allowing methods producing data on dramatically different response scales to be directly compared. We then ran a ‘*moving window*’ function in the algorithm, which iterates through all subsets in the data greater than 2 years, and subjects each ‘window’ to a linear model, which calculates the slope statistic (change of standardized density over change in year), standard error of this slope, p-value, and R^2^.

Then we ran the ‘*stability time*’ function in the same package to calculate the number of years to reach stability (Bahlai et al., 2021). The number of years to reach stability is calculated in the function by using the summary statistics previously calculated for each dataset to compute the proportion correct for each window within the standard deviation of the longest slope, and returns the number of windows whose slope are within said standard deviation (95% CI). We also ran the ‘*proportion_wrong’* function in the same package (Bahlai et al., 2021), to calculate the proportion of windows that did not reach stability with a p-value that was not within 0.05 of the p-value of the true slope (slope of the longest series). The ‘significantly wrong’ windows would likely lead a scientist to a spurious conclusion that does not agree with the overall trend. We calculated both the proportion of ‘significantly wrong’ windows before stability time and overall, for each dataset.

### Statistical analyses

Following the use of the ‘broken windows’ algorithm, we assessed if and how study parameters affected years to stability time, the overall proportion wrong, and the proportion wrong before stability was reached for each dataset. We did this by comparing the means between different groups using independent samples t-tests in base R using RStudio and R statistical software (RStudio Team, 2020; R Core Team, 2020). Using this method, we compared results from datasets that varied by life stage studied, the geographic scope of the study, sampling techniques used, and the study response variables. For example, we tested whether the mean stability time for datasets collected using dragging sampling methods was significantly different from the mean stability time for datasets collected using opportunistic sampling methods. We also used Pearson product moment correlation tests in base R using RStudio and R statistical software (RStudio Team, 2020; R Core Team, 2020). These tests were used to assess potential correlations between study length and stability time, overall proportion wrong, and the overall proportion wrong compared to proportion wrong before stability was reached. The dataset, R code, test results, and resulting figures are also provided in the Supplemental Materials.

## Results

### Study length

Using the ‘broken window’ algorithm, we found that across all datasets, years to stability time ranged from 4 to 23 years (Figure 1). The proportion of datasets that reached stability by 4 years is very low; by 10 years, the majority of datasets had reached stability (Figure 1). We found that study length was positively correlated with years to stability (Pearson product moment correlation test, r^2^ = 0.9, p-value <0.001) (Figure 2). From our analysis, we also found that the proportion of windows significantly wrong before datasets reached stability ranged from 0 to 100% (Figure 3). We also found that the association between proportion significantly wrong before stability time and overall proportion wrong were not strongly correlated (t = -1.2, df = 571.46, p-value = 0.23, Figure 3).

**Figure 1.**
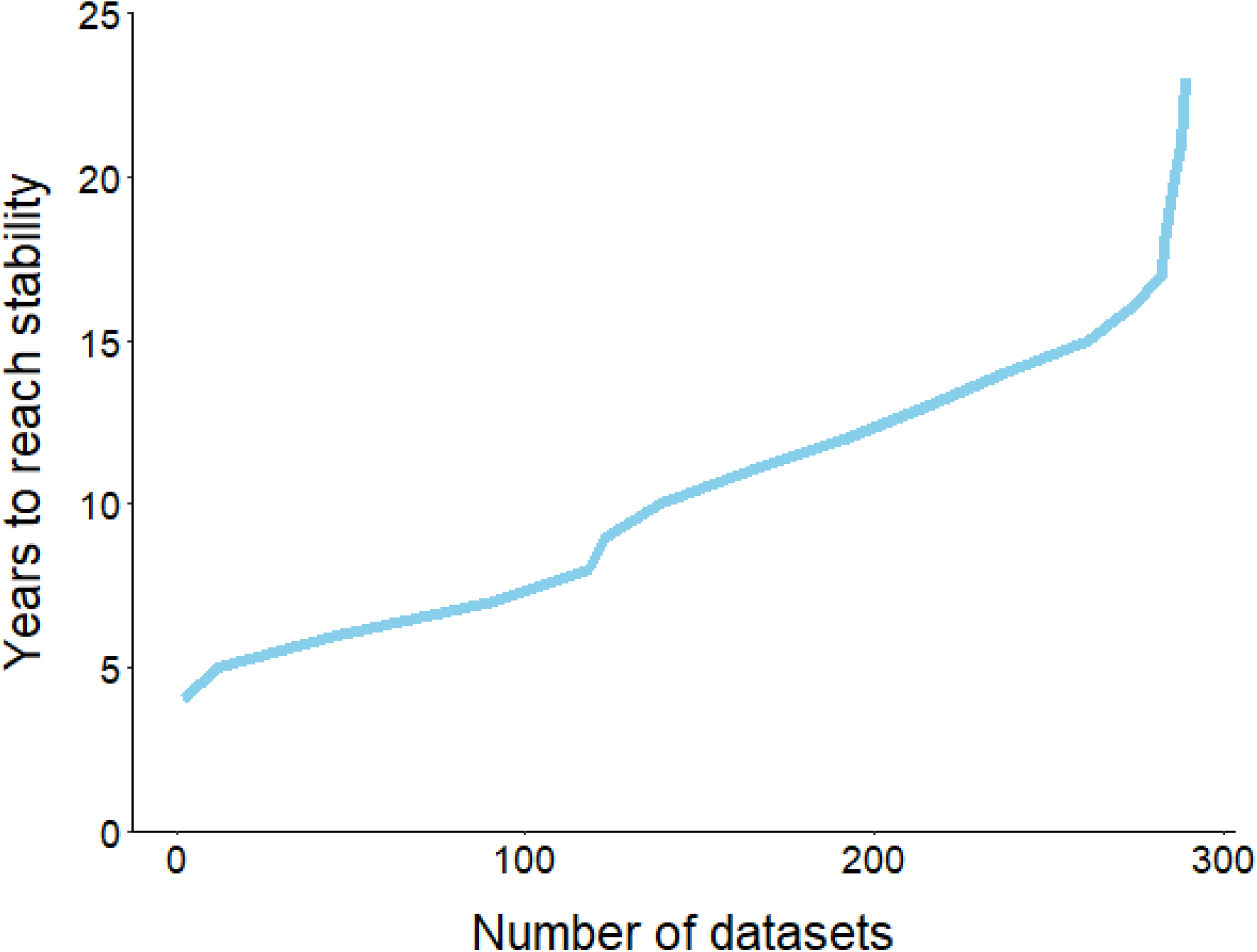
The fraction of datasets that take *y* years to reach stability. Number of years for datasets within our study to reach stability. Majority of datasets (n = 289) required 10 years to reach stability, and none reached stability before 4 years.

**Figure 2.**
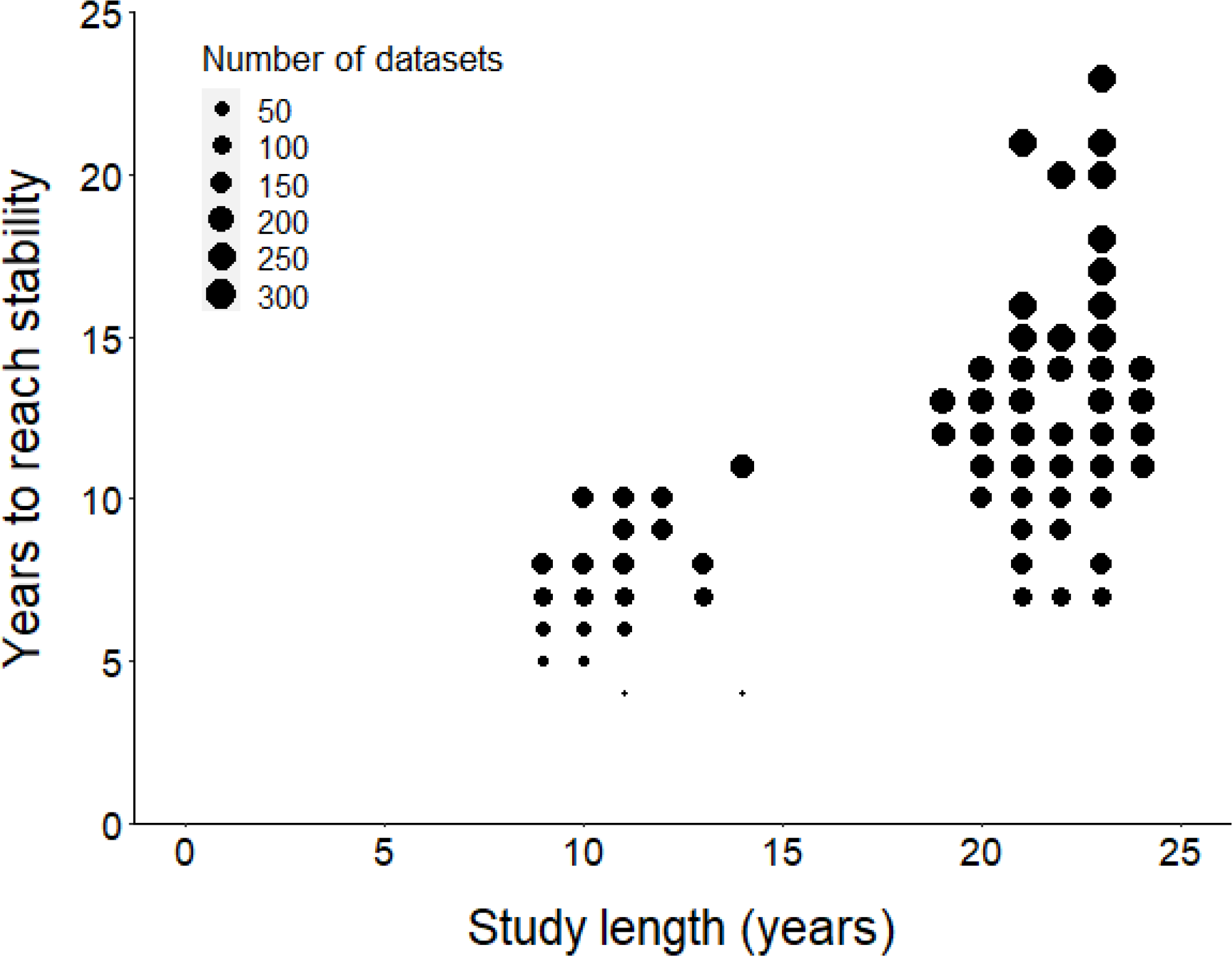
Comparison of study length, years to reach stability and the number of datasets. Scatter plot comparing study length with the number of years to reach stability. Point size represents the number of raw datasets (n=289). Overall, the number of years to reach stability increased with study length.

**Figure 3.**
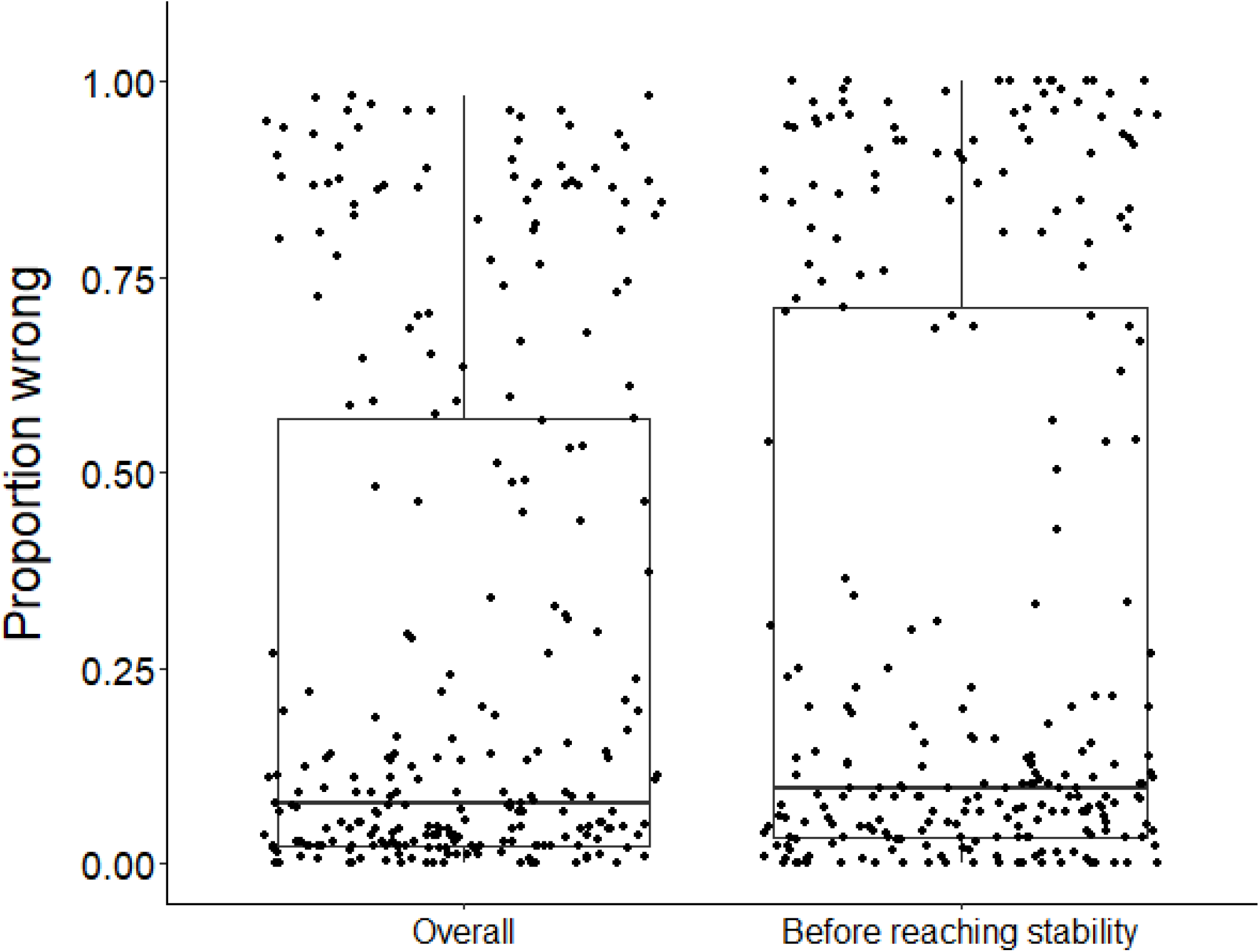
Overall proportion wrong compared to proportion wrong before stability. Whisker plot comparing the overall proportion wrong and proportion wrong before reaching stability and showing no significant differences (t = -1.2, df = 571.46, p-value = 0.23). Each dot represents the results from a dataset (n=289) analyzed.

### Sampling technique

We found that datasets with dragging techniques reached stability time faster than datasets collected using opportunistic, passive sampling methods (t = -8.53, df = 236.23, p-value <0.001) (Figure 4A). The median number of years to reach stability was 7 for dragging techniques and 12 for opportunistic techniques. However, we did not find a significant difference when we compared the proportion significantly wrong before stability by sampling technique (t = 0.08, df = 155.58, p-value = 0.93, Figure 4B). Datasets with dragging sampling techniques in our data experiment also exhibited a smaller range of years to stability, compared to datasets with opportunistic passive sampling methods.

**Figure 4.**
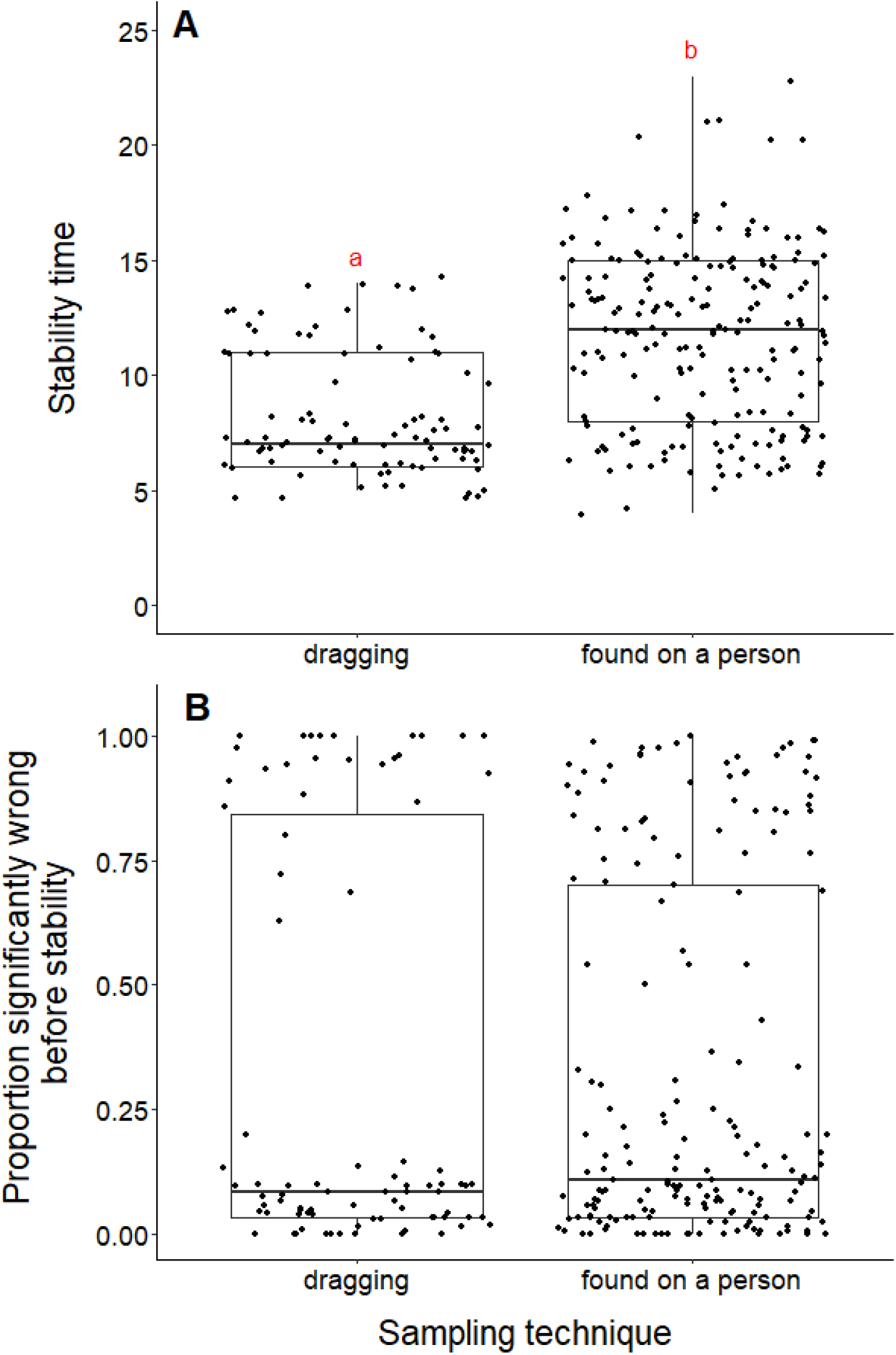
Comparison of stability time and proportion significantly wrong before stability by sampling techniques. A) Years to reach stability by sampling technique, dragging (n=90) and public survey (n=198). Letters above boxes denote significant differences (t=-8.53, df=236.23, p-value < 0.001). B) Proportion significantly wrong before reaching stability across sampling technique, showing no significant difference between techniques (t=0.08, df = 155.58, p-value = 0.93).

### Life stage

Datasets measuring black-legged tick larvae reached stability slower than datasets measuring adult or nymph abundance data (adults and larvae, t = -5.96, df = 10.11, p-value <0.001; nymphs and larvae, t = -5.52, df = 10.54, p-value <0.001, Figure 5A). We found no difference in the time to stability between datasets measuring adults or nymph blacklegged ticks (t = -0.64, df = 126.84, p-value = 0.52). The median time for datasets measuring larvae to reach stability was 11.5 years, compared to 7 years for adults and nymphs.

**Figure 5.**
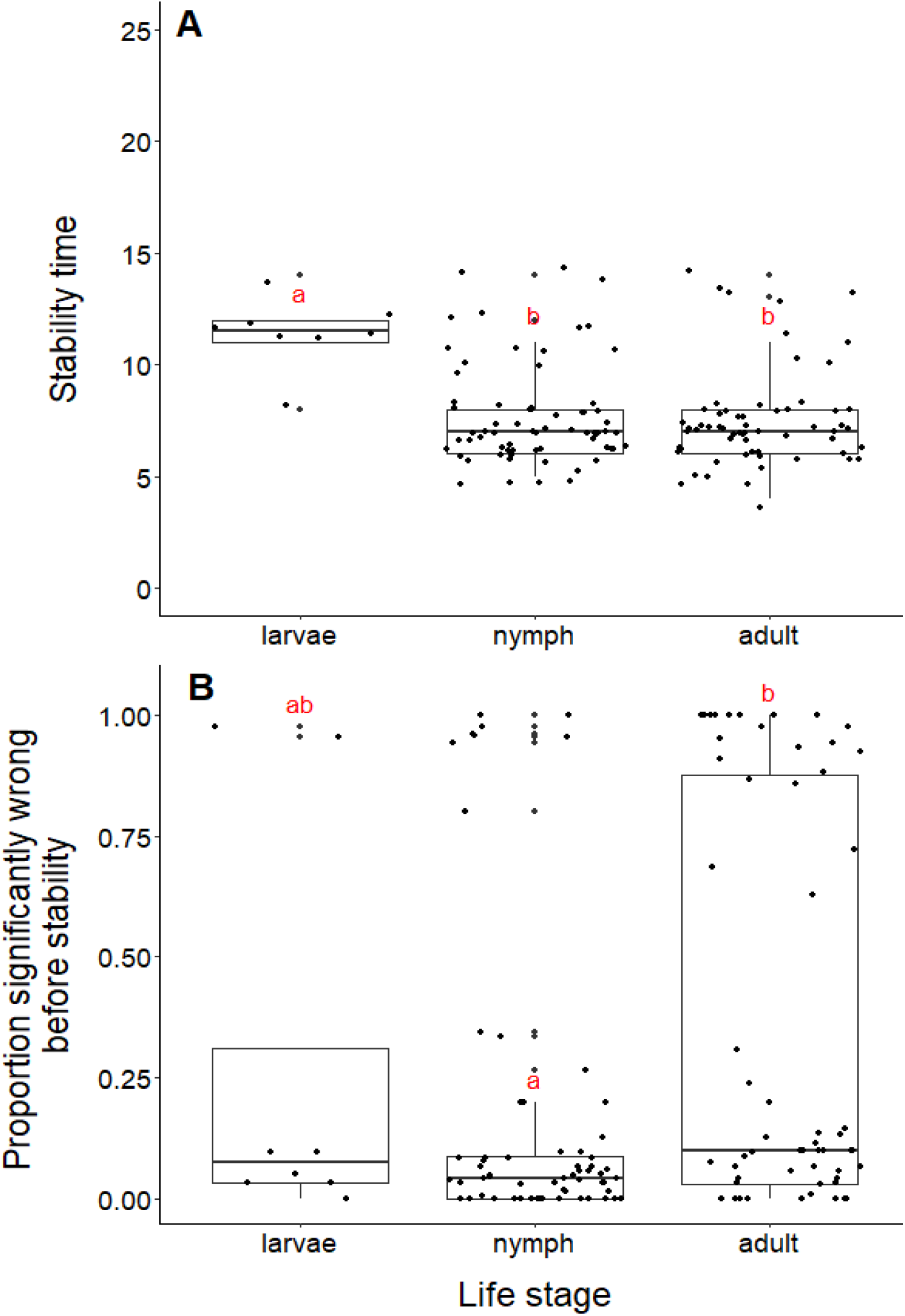
Comparison of stability time and proportion significantly wrong before stability by life stage. A) Years to reach stability by life stage, larvae (n=8), nymphs (n=68), and adult (n=63). Letters above boxes denote significant differences between larvae and adults (t = -5.96, df = 10.11, p-value <0.001) and larvae and nymphs (t = -5.56, df = 10.33, p-value <0.001), while there is no significant difference between nymphs and adults (t = -0.63, df = 128.99, p-value = 0.53). B) Proportion significantly wrong before reaching stability across life stage. Letters above boxes denote significant differences between nymphs and adults (t = 2.99, df = 112.55, p-value = 0.003). There is no significant differences between nymphs and larvae (t = -0.78, df = 7.85, p-value = 0.46) or adults and larvae (t = 0.44, df = 8.77, p-value = 0.67).

The mean proportion wrong before stability for adults was 0.35, compared to nymphs (0.16) and larvae (0.28). Datasets measuring nymph tick life stages had a significantly lower proportion of windows wrong before stability time compared to adults (t = 2.90, df = 113.64, p-value = 0.004, Figure 5B). We found no difference in the proportion wrong before stability time between larvae and nymphs (t = -0.75, df = 7.89, p-value = 0.47) and larvae and adults (t = 0.44, df = 8.77, p-value = 0.67)

### Geographic scope

Blacklegged tick datasets from studies reporting data from counties reached stability in fewer years than datasets reported at smaller spatial scales (town, state forest, or grid) (county and grid, t = -17.21, df = 33.87, p-value <0.001; county and state forest, t = -5.55, df = 5.22, p-value = 0.002; county and town, t = -17.03, df = 243.09, p-value <0.001; Figure 6A). The median number years to stability for datasets of county scope was 7 years, compared to 11.5 years for grid and 12 years for town and state forest. We found no difference in years to stability comparing across datasets reported from towns, state forests, or grids (town and state forest, t = 0.47, df = 6.12, p-value = 0.65; town and grid, t = 0.22, df = 86.18, p-value = 0.82; state forest and grid, t = -0.38, df = 6.03, p-value = 0.72; Figure 6A).

**Figure 6.**
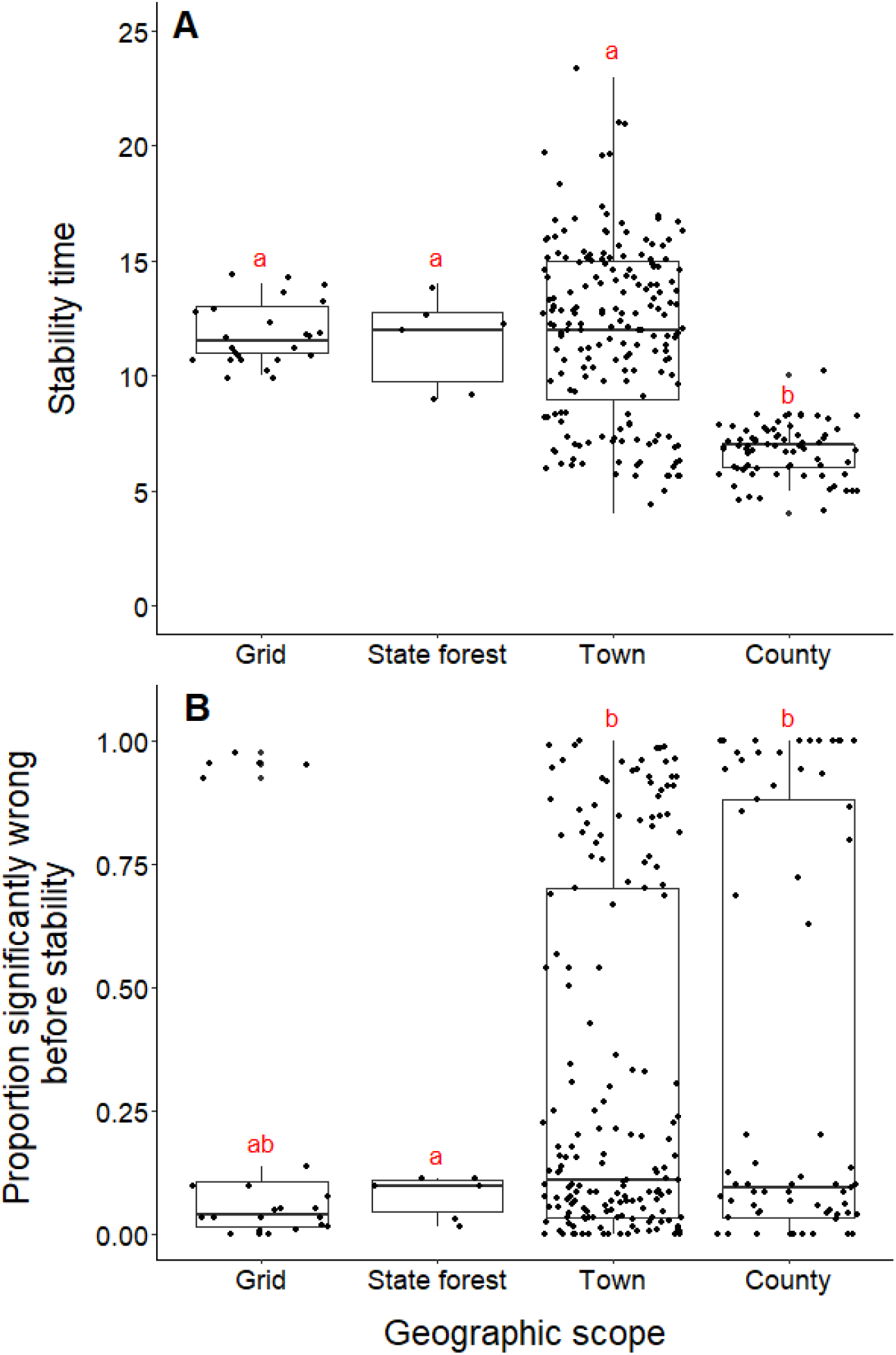
Comparison of stability time and proportion significantly wrong before stability by geographic scope. A) Years to reach stability by geographic scope, grid (n=24), state forest (n=6), town (n=186) and county (n=73). Letters above boxes denote significant differences between county and town level studies (t = - 17.03, df = 243.09, p-value <0.001); county and state forest level studies (t = -5.55, df = 5.22, p-value = 0.002); county and grid level studies (t = -17.21, df = 33.87, p-value <0.001). There is no significant between town and state forest level studies (t = 0.47, df = 6.12, p-value = 0.65); town and grid level studies (t = 0.22, df = 86.18, p-value = 0.8); state forest and grid level studies (t = -0.38, df = 6.03, p-value = 0.72). B) Proportion significantly wrong before reaching stability across geographic scope. Letters above boxes denote significant differences between state forest and county level studies (t = 5.24, df = 74.29, p-value <0.001); state forest and town level studies (t = 7.42, df = 46.44, p-value <0.001). There is no significant difference between county and town level studies (t = 0.72, df = 115.03, p-value = 0.48); county and grid level studies (t = 1.33, df = 43.07, p-value = 0.19); town and grid level studies (t = 1.00, df = 28.52, p-value = 0.32); state forest and grid level studies (t = -1.91, df = 25.10, p-value = 0.07).

Datasets from studies reporting data from state forests had a significantly lower proportion of wrong windows, as measured by the proportion wrong before stability time, compared to datasets reported from counties and towns (state forest and county, t = 5.24, df = 74.29, p-value <0.001; state forest and town, t = 7.42, df = 46.44, p-value = p-value <0.001; Figure 6B). Studies reported from grids did not have a higher proportion of windows wrong before stability time (t = -1.91, df = 25.10, p-value = 0.068). The mean proportion significantly wrong for data at the scale of state forest is 0.077, compared to 0.35 of county datasets, 0.31 of town datasets, and 0.23 of grid datasets. The proportion of wrong windows did not significantly vary between county, town, and grid studies (county and town, t = 0.72, df = 115.03, p-value = 0.48; county and grid, t= 1.33, df = 43.07, p-value = 0.19; town and grid, t = 1.00, df = 28.52, p-value = 0.32; Figure 6B).

### Sampling metric

We found no difference in the number of years until a dataset reached stability time comparing datasets measuring the abundance of black-legged ticks and datasets that tested for *B. burgdorferi* infection (t = -1.29, df = 283.9, p-value = 0.20; Figure 7A). The datasets also did not significantly vary in the proportion of wrong windows, as measured by the proportion wrong before stability time (t = -1.18, df = 232.98, p-value = 0.24; Figure 7B).

**Figure 7.**
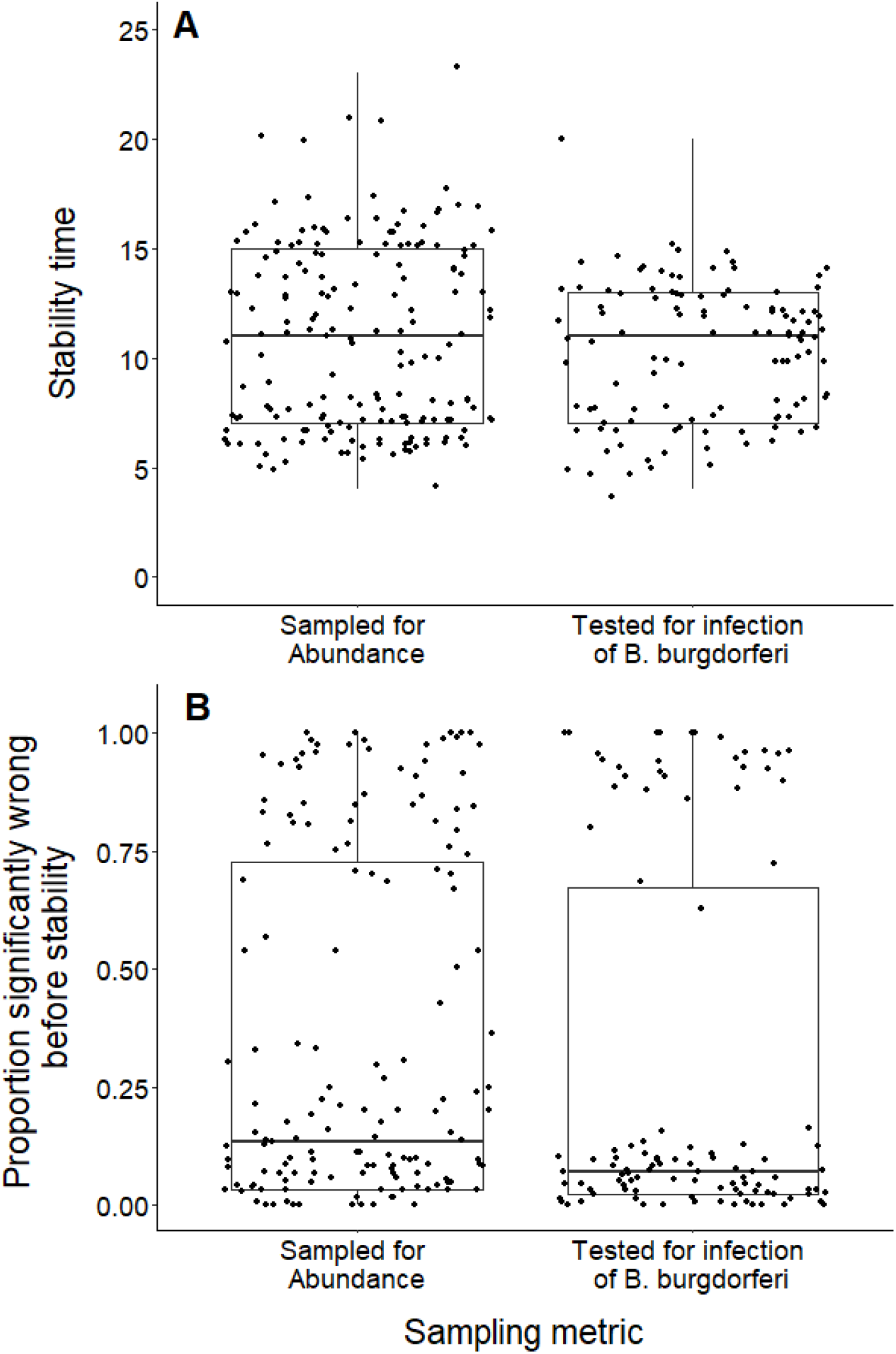
Comparison of stability time and proportion significantly wrong before stability by study response variable. A) Years to reach stability by study response variable, deer tick abundance (n=175) and *B. burgdorferi* presence in deer ticks (n=114) and showing no significant difference between study response variables (t = -1.29, df = 283.9, p-value = 0.20). B) Proportion significantly wrong before reaching stability across study response variables and showing no significant difference between study response variables (t = - 1.18, df = 232.98, p-value = 0.24).

## Discussion

Overall, we found several aspects of how blacklegged ticks are monitored and data reported may have an impact on the likelihood of a study reflecting longer-term population trends. This is supported by our findings about datasets reaching stability time, as well the proportion of significantly wrong windows, varying by study methods. Specifically, datasets using dragging, counting nymph or adult life stages, and reported by county, reached stability faster than other datasets. These findings support our hypothesis that the dragging method would be associated with more consistent patterns than other sampling methods. We also predicted that datasets collected on different blacklegged tick life stages may vary in stability time; however, we did not anticipate that larvae would reach stability time slower than other life stages. Our results also indicate the importance of long term monitoring, as we found that the longer datasets were correlated with more years to reach stability and many datasets with higher proportions of misleading windows before stability time. As none of the datasets assessed reached stability time before 4 years, our analyses indicate that short term studies may be insufficient to explain long term dynamics in blacklegged ticks.

Our findings are consistent with previous research supporting the importance of long term ecological monitoring for understanding populations in field biology. Several other studies provide support for using long term studies to understand patterns (White, 2019; Cusser et al., 2020; Cusser et al. 2021). For example, White found that at least 72% of the vertebrate populations they studied required at least 10 years of study to detect a significant trend in abundance, while Cusser et al. determined that it would take at least 16-19 years to detect a significant effect from no-till management on crop yield and soil moisture, emphasizing the importance of long term studies. Similarly, we predicted and found that the dragging sampling technique would produce more consistent abundance trends than datasets associated with opportunistic surveying. This was expected given research indicating that dragging can be a consistent way to monitor black-legged ticks (Falco and Fish, 1992). However, other research has found that data produced using different tick sampling methods is similar (Rulison et al., 2013). Yet even more recent research attempting to validate dragging with other tick species in Sweden concluded that dragging was not recommended for tick adults or larvae, due to low repeatability and agreement (Kjellander et al., 2021). This may be due to the lack of standardization within dragging cloths or other within-method variability between surveys (Newman et al., 2019).

We anticipated, and found, that datasets studying different life stages of ticks would vary in how many years it took to reach stability time. Specifically, time to stability was longer for datasets monitoring larvae, the youngest life stage, compared to adults and nymphs. Additionally, datasets monitoring adult black-legged ticks had higher proportions of significantly wrong windows. This finding is critical for researchers trying to glean insights into broad scale black-legged tick trends across disparate study methods. This suggests that datasets that focus on adults will be more likely to lead to misleading results if the study has not reached stability, which supports longer-term studies. This finding is critical for understanding how to apply black-legged tick monitoring data to potential Lyme disease risk, as different life stages may pose different levels of exposure risk to humans. Nymph, not adult, abundance has been shown to be the primary factor in determining risk of exposure (Falco et al., 1999). Due to this potential variation in transmission by life stage, the recent US Centers for Disease Control and Prevention (CDC) report on black-legged tick surveillance provides distinct guidance on collecting and interpreting data from each life stage (CDC 2021). Our results support this distinction. However, life stage is distinct from other categorizations of tick life history thought to impact abundance and disease transmission, such as ‘physiological age,’ a measure that groups ticks based on environmental stress events (Pool et al., 2017). There may be important variation among ticks of the same life stage, based on their physiological age, that surveyors are not assessing in the field, as this may require the addition of molecular biology approaches (Estrada-Peña et al., 2021). And even though we found no difference in our results when comparing the number of years to reach stability and proportion wrong before stability in datasets that measured black-legged tick abundance and the percent with *Borrelia burgdorferi*, this may be due to additional variation within studies measuring pathogen prevalence within ticks. Some studies test pools of ticks for infection to reduce cost and labor, and variation in the number of ticks processed can impact the perceived estimation of prevalence (Estrada-Peña et al., 2021).

Our results indicate that sampling a larger geographic area leads to more consistent patterns in black-legged tick population dynamics than studies conducted on smaller areas. This is supported by previous work finding high levels of fine-scale spatial distribution of black-legged ticks (Dumas et al., 2022, Mathews-Martin et al., 2020). Fortunately, county-level information is a common geographic scale for documenting the presence and abundance of blacklegged ticks and Lyme disease (e.g. Fleshman et al., 2021). Given that ecological monitoring requires tradeoffs in effort, cost, and insight (Bennett et al., 2018), this finding may indicate that studies at larger spatial scales may be able to give indication of population trajectories faster than studies at smaller scales.

Our findings also support the collection of long term tick datasets. Even with the improved insights from longer monitoring studies, managers and policymakers will still need to make decisions without complete information. Determining what is ‘enough’ data for evidence-based decision making is a challenge (Canessa et al., 2015). As the lack of shared, standardized sampling methods may hamper the ability to monitor changes in abundance and distribution of ticks (Dennis et al., 1998; Eisen et al., 2016), understanding patterns and differences between datasets may help. Our results can help inform if and how black-legged tick monitoring data can be reasonably interpreted based on study parameters to make informed, rigorous claims to support management decisions that can help reduce Lyme disease exposure and risk. Moreover, we further demonstrate the value of resampling existing data to give insights into analyzing existing data and designing future monitoring programs (White and Bahlai, 2021).

Understanding population trajectories is also critical for using ecological data for ecological forecasting. Ecological forecasting is considered the next critical frontier in broad scale environmental science (Dietze et al., 2018). Ecological forecasting initiatives are increasing, with an NSF-funded Research Coordination Network (NSF, 2019), conferences (EFI, 2019; EFI RCN and NEON, 2020), and ongoing ecological forecast challenges (EFI RCN NEON, 2020). With study organisms such as black-legged ticks, forecasting could be used to proactively protect human health. As standardized data has been identified as critical for understanding tick-transmitted pathogens (Estrada-Peña et al., 2021), future work should examine how these study parameters, identified as critical through retrospective data resampling, shape forecasted patterns and simulations. This is especially important for forecasting disease risk, as data gaps and variation in monitoring data are already considered major challenges (Kugeler and Eisen 2020). Our results highlight the need to better understand and explicitly incorporate that variability into the interpretation of black-legged tick data.

## Conclusions

We used 289 datasets from 7 long term black-legged tick studies to conduct a moving windows data resampling experiment, revealing that sampling design does shape inferences in black-legged tick population trajectories and the number of years to reveal stable patterns. Specifically, our results show the importance of study length, sampling technique, life stage, and geographic scope in understanding black-legged tick populations, in the absence of standardized surveillance methods. Current public health efforts based on existing black-legged tick datasets must take monitoring study parameters into account, to better understand if and how to use monitoring data to inform decisioning. We also advocate that potential future forecasting initiatives consider these parameters when projecting future black-legged tick population trends. Understanding how study design and monitoring shapes synthesis is critical across many biological disciplines.

## Acknowledgments

RMC was supported by the National Institute of General Medical Sciences of the National Institutes of Health under Award Number R25GM122672. CAB, JP, and KSW were supported by the Office of Advanced Cyberinfrastructure in the National Science Foundation under Award Number #1838807. The content is solely the responsibility of the authors and does not necessarily represent the official views of the National Institutes of Health or the National Science Foundation.

## Supplemental Materials

The data for this study are located here -

Rowan Christie, Kaitlin Stack Whitney, Julia Perrone, & Christie Bahlai. (2022). Ixodes Scapularis monitoring data compiled from 7 studies [Data set]. Zenodo.

https://doi.org/10.5281/zenodo.6540831

The code for this study is located here:

Rowan Christie. (2022). SMCCoder/ixodes_scapularis_research: Ixodes scapularis research code for PeerJ submission (v1.1). Zenodo. https://doi.org/10.5281/zenodo.6820316

### Appendix 1. PRISMA workflow for finding and screening black-legged tick studies

Screening criteria: We searched for datasets that were at least 9 years and measured black-legged tick density or abundance using any sampling method, documenting the process and results using the PRISMA flow diagram (Moher et al., 2009).

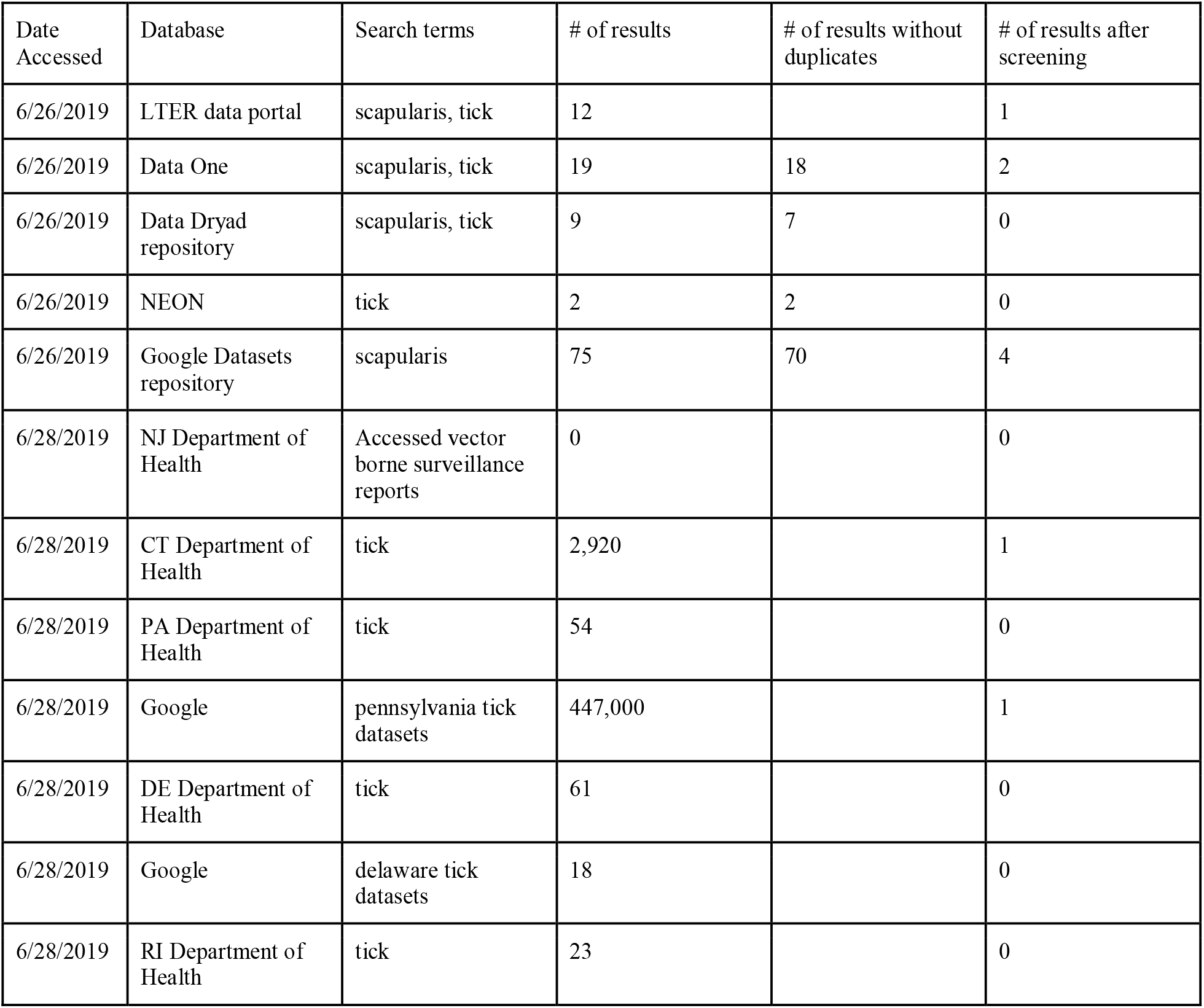

