## Appendix 1 for "Longer study length, standardized sampling techniques, and broader geographic scope leads to higher likelihood of detecting stable abundance patterns in long term black-legged tick studies"

**Appendix 1.  PRISMA workflow for finding and screening black-legged tick studies**

Screening criteria: We searched for datasets that were at least 9 years and measured black-legged tick density or abundance using any sampling method, documenting the process and results using the PRISMA flow diagram (Moher et al., 2009).

| Date Accessed | Database | Search terms | # of results | # of results without duplicates | # of results after screening |
| --- | --- | --- | --- | --- | --- |
| 6/26/2019 | LTER data portal | scapularis, tick | 12 |  | 1 |
| 6/26/2019 | Data One | scapularis, tick | 19 | 18 | 2 |
| 6/26/2019 | Data Dryad repository | scapularis, tick | 9 | 7 | 0 |
| 6/26/2019 | NEON | tick | 2 | 2 | 0 |
| 6/26/2019 | Google Datasets repository | scapularis | 75 | 70 | 4 |
| 6/28/2019 | NJ Department of Health | Accessed vector borne surveillance reports | 0 |  | 0 |
| 6/28/2019 | CT Department of Health | tick | 2,920 |  | 1 |
| 6/28/2019 | PA Department of Health | tick | 54 |  | 0 |
| 6/28/2019 | Google | pennsylvania tick datasets | 447,000 |  | 1 |
| 6/28/2019 | DE Department of Health | tick | 61 |  | 0 |
| 6/28/2019 | Google | delaware tick datasets | 18 |  | 0 |
| 6/28/2019 | RI Department of Health | tick | 23 |  | 0 |
